# Labelling effect in insects: cue associations influence perceived food value in ants

**DOI:** 10.1101/680959

**Authors:** Stephanie Wendt, Tomer J. Czaczkes

**Affiliations:** Animal Comparative Economics laboratory, Institute of Zoology & Evolutionary Biology, University of Regensburg, 93053 Regensburg, Germany

**Keywords:** Product labels, Associative learning, Relative value perception, Assimilation, Branding

## Abstract

Humans usually assess things not in terms of absolute value, but relative to reference points. The framing of alternatives can strongly affect human decision-making, leading to different choices depending on the context within which options are presented. Similar reference-point effects have been recently reported in ants, in which foragers show a contrast effect: ants overvalue a medium-quality food source if they were expecting a poor one, and vice versa for expectations of good food. However, studies of human consumer psychology have demonstrated that expectations, for example from product labels, can drive value perception in the other direction via assimilation. For example, an expensive bottle of wine is perceived as more enjoyable compared to a cheaper bottle, even if the wine is the same. In this study, we demonstrate a similar labelling-association effect in an insect: ants showed assimilation effects by spending twice as long drinking at a medium quality food source if it was scented with an odour previously associated with high quality than if it was scented with a poor-quality label. The presence of odour cues in the food during consumption and evaluation is critical, as without them, odour-driven expectations of quality result in contrast, not assimilation effects. The addition of a quality label in the food thus reverses contrast effects and causes value to be aligned with expectations, rather than being contrasted against them. As value judgement is a key element in decision-making, relative value perception strongly influences which option is chosen, and ultimately how choices are made.

## Introduction

A decision is often made by evaluating and comparing the available options, which then usually leads to a choice for the option promising the greatest profit (von Neumann and Morgenstern 1944). The way in which options are evaluated may, however, strongly influence which option is ultimately chosen. Thus, understanding the factors influencing the perceived value of available options helps us to understand human behaviour and decision making (Slovic 1995; Thaler and Sunstein 2008; Tversky and Kahneman 1981). Understanding the drivers of option evaluation and comparison are thus central to the study of behavioural economics and consumer psychology.

Although early economic theories described humans as rational decision-makers who always choose the option with the greatest utility regardless of other factors (von Neumann and Morgenstern 1944; Vlaev et al. 2011), a large body of evidence has accumulated demonstrating that this is not always the case. Kahneman and Tversky (1979) suggested that decision-making is not based on absolute outcomes, but rather on the relative perceptions of gains and losses. According to Prospect Theory, the value of options being evaluated is determined relative to a reference point, such as the status quo or former experience (Kahneman and Tversky 1979; Parducci 1984; Tversky and Kahneman 1992; Ungemach, Stewart, and Reimers 2011; Vlaev et al. 2011). Thus, an option which leads to a loss compared to the reference point is perceived more negatively than if the same option led to a gain compared to the reference (Kahneman and Tversky 1979). For example, satisfaction gained from income is perceived not absolutely, but relative to the income of one’s colleagues (Boyce, Brown, and Moore 2010). Therefore, human decision making tends to be relative rather than rational.

The concept of malleable value perception is not just relevant to humans. Value judgments in animals are also influenced by factors apparently independent of the absolute value of options, such as the state an animal is in during learning (Aw et al. 2009; Aw, Vasconcelos, and Kacelnik 2011; Czaczkes, Brandstetter, et al. 2018; Lydall, Gilmour, and Dwyer 2010; Pompilio, Kacelnik, and Behmer 2006) and expectations (Annicchiarico et al. 2016; Bentosela et al. 2009; Bitterman 1976; Couvillon and Bitterman 1984; Crespi 1942; Flaherty 1982, 1999; Mustaca, Bentosela, and Papini 2000; Oberhauser and Czaczkes 2018; Papini et al. 2001; Flavio Roces and Núñez 1993; Flavio Roces 1993; Webber et al. 2015; Weinstein 1970; Wendt et al. 2019). Expectations can cause identical options to be perceived differently depending on whether a better or worse option was expected instead of the presented one. In a previous study, for example, ants showed lower food acceptance towards medium quality food when they expected high quality food (negative contrast) and higher acceptance of medium food when expecting poor food (positive contrast) (Wendt et al. 2019). Similarly, capuchin monkeys refuse otherwise acceptable pay (cucumber) in exchanges with a human experimenter if they had witnessed a conspecific obtain a more attractive reward (grape) for equal effort (Brosnan and de Waal 2003). Bees too rejected otherwise acceptable lower quality food when they expected high quality food due to previous experience (Bitterman 1976; Couvillon and Bitterman 1984). Such incentive contrast effects (Flaherty 1999) represent one of the main influences on subjective value. We see incentive contrasts as a subset of relative value perception.

In humans, product and brand labels alter perceived value, and directly affect purchasing decisions. Such labels convey expectations, and thus reference points for judging an option (French and Smith 2013). Depending on previous associations with the label, perceived option value can increase (Breneiser and Allen 2011; Fornerino and d’Hauteville 2010; Kühn and Gallinat 2013; Lee et al. 2013; Nevid 1981; McClure et al. 2004; Wansink 2000; Woodside and Taylor 1978; Yamada et al. 2014) or decrease (Lee, Frederick, and Ariely 2006; Wansink 2000). For example, drinks presented along with strong brands such as “Coca Cola” (which have strong positive associations due to successful marketing campaigns) tend to be rated as being tastier or more attractive compared to identical drinks which were presented with weaker brand labels or without any labels, even though there is rarely a preference found in blind tests (Breneiser and Allen 2011; Fornerino and d’Hauteville 2010; Kühn and Gallinat 2013; McClure et al. 2004; Yamada et al. 2014). Compared to these strong international brands, store brands are often believed to offer lower product quality and nutritional value (Cunningham, Hardy, and Imperia 1982; Dick, Richardson, and Jain 1995). If the difference between a label-driven expectation and the objective value is small, the perceived value aligns with the expectation in a process called assimilation (Cardello and Sawyer 1992; Hovland, Sherif, and Harvey 1957; Schnurr, Brunner-Sperdin, and Stokburger-Sauer 2017). For example, a cola drink which previously received a low rating may receive a significantly better rating when subjects were told that it is of a favourable brand (Cardello and Sawyer 1992). In humans, labels are an accumulation of various associative cues which evoke a positive or negative response once the label is seen (French and Smith 2013; Macklin 1996). Such associated attributes may affect value perception in animals as well.

Associative learning, through which cues or actions are learned to predict a positive or negative experience, is almost ubiquitous in the animal kingdom (Couvillon and Bitterman 1980; Giurfa 2007; Hawkins and Byrne 2015; Menzel 1993; Rankin 2004; Nelson, Oxley, and Clawson 1997; Sahley, Rudy, and Gelperin 1981; Siwicki and Ladewski 2003; Spatz, Emanns, and Reichert 1974). Like associative labelling in humans, perceived option value varies for animals as well. Naïve ants, for example, prefer food presented alongside an odour which had already been received through food exchanges inside the nest over food presented with a novel odour, because the familiar odour was previously associated with a positive event (Provecho and Josens 2009). An example for negative associations was shown in leaf cutter ants: Odour cues associated with damage to the ants’ cultivated fungus drive aversion to otherwise acceptable fungal substrate, with the odour cue acting as a negative food label (Roces 1994; Saverschek and Roces 2011).

The aim of this study was to investigate whether labelling effects as shown in humans can be demonstrated in insects, and whether it is affected by the point of time of cue presentation. We thus ask whether ants align their perception of a food sources’ value with value-associated odour cues presented in the food during consumption. We previously demonstrated a contrast effect in ants, whereby ants undervalue or overvalue food if they were expecting something of better or worse quality, respectively (Wendt et al. 2019). In that case, expectations generated *before* perception of the objective food quality drove value perception. Here, we ask how value-related labels experienced *during* consumption affect perceived value. We offered ants food which would, due to negative contrast effects, normally be rejected, and presented it along with previously positively or negatively associated odour cues. We hypothesized that incentive contrast effects could be counteracted by the mere presence of associative odour cues during consumption.

## Methods

### Study animals

Eight stock colonies of the black garden ant *Lasius niger* were collected on the University of Regensburg campus. The colonies were housed in 30×30×10cm foraging boxes with a layer of plaster covering the bottom. Each box contained a circular plaster nest box (14 cm diameter, 2 cm height). The colonies were queenless with around 1000-2000 workers and small amounts of brood. Queenless colonies forage and lay pheromone trails, and are frequently used in foraging experiments (Devigne and Detrain 2002; Dussutour et al. 2004). The colonies were fed with 0.5M sucrose solution and received *Drosophila* fruit flies once a week. Colonies were deprived of sucrose solution four days prior to the experiments in order to achieve a uniform and high motivation for foraging (Mailleux, Detrain, and Deneubourg 2006; Josens and Roces 2000). Water was always available *ad libitum*.

### General setup

The setup consisted of a 20 × 1 cm long paper-covered runway which was connected to the colony’s nest box via a 40 cm long drawbridge (figure 1A). A 5mm diameter drop of sucrose solution (Sigma-Aldrich) was placed on an acetate feeder (2 × 1.5 cm) at the end of the runway (60cm from the nest). The molarity of the sucrose droplet depended on the experiment, treatment, and on the ants’ number of visit to the food source.

**Fig. 1:**
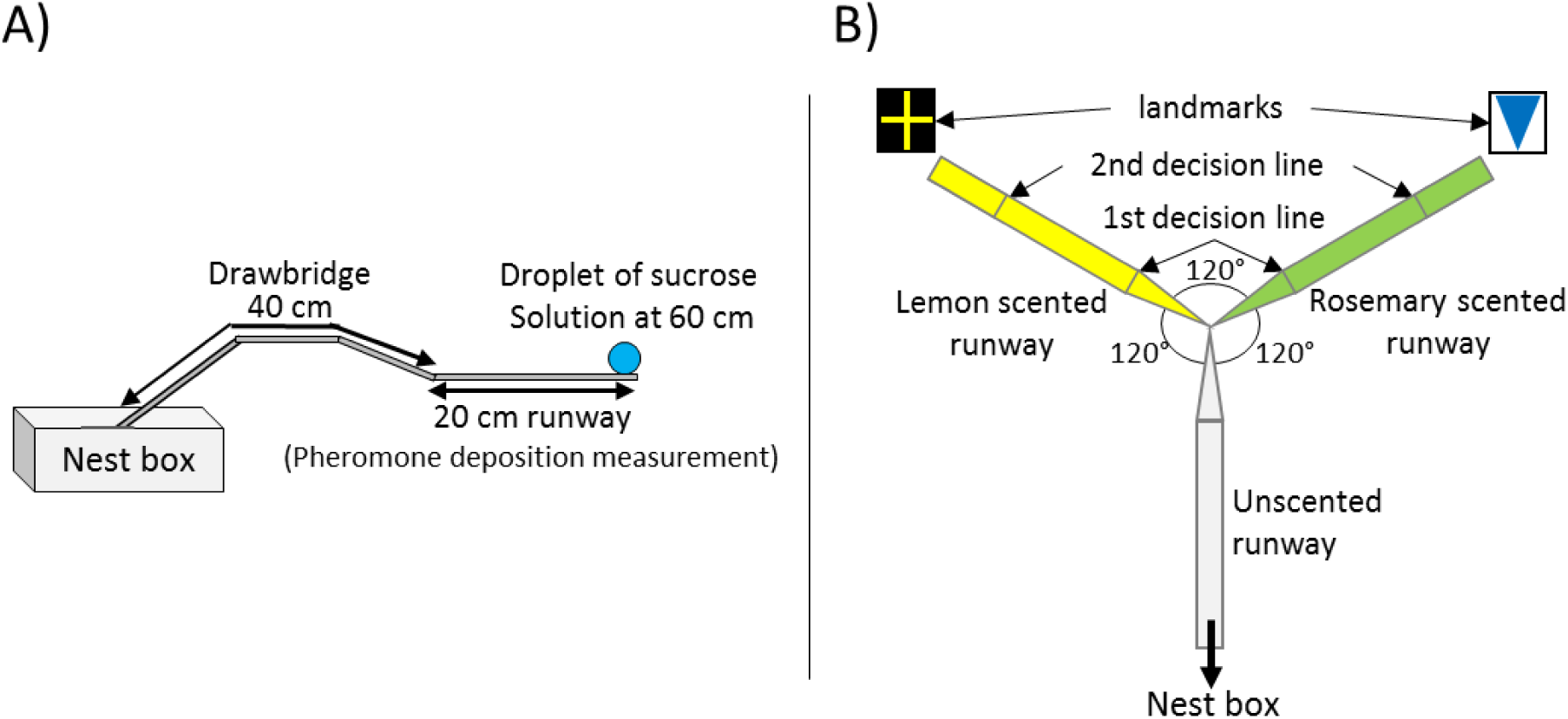
**A)** General setup used for all presented experiments. The 20 cm long runway is connected to the nestbox via a 40 cm long drawbridge. The droplet of sucrose solution is placed at the end of the runway (60 cm distance to the nest). All runways were covered with paper overlays while the overlay on the last bit was replaced by pheromone-free ones each time an ant had walked over it. **B)** Y-maze used on the 10^th^ visit of experiment 2. All arms were 10 cm long. The arm connected to the nest box was covered with unscented paper overlays while the other two arms were covered with lemon and rosemary scented paper overlays (one scent on each side). Visual cues (landmarks) were placed right behind the two scented arms. The first decision line was located 2.5cm from the y-maze centre and marked the initial decision of an ant while the second decision line was placed 7.5cm from the centre and marked the final decision.

To begin an experiment, the sub-colony was connected to the runway via the drawbridge. 2-4 ants were allowed onto the runway, and the first ant to reach the feeder was marked with a dot of acrylic paint on its abdomen. The marked ant was allowed to drink to repletion at the food source, while all other ants were returned to the nest. As the ant drank at the droplet it was given one of three food acceptance scores, following Wendt et al. (2019). Full acceptance (1) was scored when the ant remained in contact with the drop from the moment of contact and did not interrupt drinking within 3 seconds of initial contact. Partial acceptance (0.5) was scored if feeding was interrupted within 3 seconds after the first contact with the food source, but the ant still drank to satiety (filled its crop, as can be seen by the distention of the abdominal tergites) within 10 minutes. Lastly, rejection (0) was scored if the ant refused to feed at the sucrose solution and either returned to the nest immediately or failed to fill its crop within 10 minutes. In addition to measuring food acceptance, we also measured the time it took until the ant interrupted drinking for the first time, and the total drinking time.

When the ant had drunk to satiety or decided not to feed at the sucrose droplet, it was allowed to move back into the nest. Inside the nest, the ant unloaded the collected sugar load to its nestmates and was then allowed back onto the runway for another visit. The drawbridge was now used to selectively allow only the marked ant onto the runway. As an additional measure of perceive value, we counted the pheromone depositions the ant performed on the way to and from the feeder. Individual pheromone deposition behaviour correlates with the (perceived) quality of a food source (Beckers, Deneubourg, and Goss 1993; Czaczkes, Koch, et al. 2018; Hangartner 1970; Wendt et al. 2019). Individual ants can adapt the strength of a pheromone trail by either depositing pheromone or not, or varying the intensity of pheromone depositions (Hangartner 1970; Beckers, Deneubourg, and Goss 1993). Pheromone deposition behaviour in *Lasius niger* is highly stereotypic. To deposit pheromone, an ant briefly interrupts running to bend its abdomen and press the tip onto the ground (Beckers, Deneubourg, and Goss 1992). This allows the strength of a pheromone trail to be quantified by counting the number of pheromone depositions over the 20 cm runway leading to the feeder. Pheromone depositions were measured each time the ant moved from the food source back to the nest (inward trip), and each time the ant moved from the nest towards the food source (outward trip). Because *Lasius niger* foragers almost never lay pheromone when they are not aware of a food source (Beckers, Deneubourg, and Goss 1992), we did not measure pheromone depositions for the very first outward trip (visit 1). The presence of trail pheromone on a path depresses further pheromone deposition (Czaczkes et al. 2013). Thus, each time an ant had passed the 20 cm runway, the paper overlay covering the runway was replaced by a fresh one. All experimental runs were recorded with a Panasonic DMC-FZ1000 camera to allow for later video analysis. After each experimental run the ant was permanently removed from the colony.

## Experimental Procedure

### Overview

We trained ants to associate a high sucrose molarity (1.5M) with one scent, and a low molarity (0.1M) with a different scent. Then, in the testing phase, we used scents on the runway to trigger an expectation of either high or low molarity, which was then contrasted with a medium (0.387M) solution containing one of the learned odour cues. The molarity of the medium quality solution (0.387M) was chosen because we wanted to present the ants with identical relative increases in sucrose molarity. 0.1M was chosen as the low food quality as it is suggested to be the minimum sucrose concentration which *L. niger* ants reliably detect, distinguish, and accept (Detrain and Prieur 2014). 1.5M was chosen as the high food quality as acceptance scores plateau after 1.5M (Wendt et al. 2019).

### Detailed methods

Training to associate food quality with odour cues took place over 8 visits. The quality of the sucrose solution offered at the end of the runway alternated each visit, always beginning with the low quality solution. The solutions were scented using either rosemary or lemon essential oils (0.5µl essential oil per ml sucrose solution, rosemary: *Rosmarinus officinalis*; Lemon: *Citrus limon*, Markl GbR, Grünwald). In half the trials the 1.5M solution was scented with lemon and the 0.1M with rosemary, and vice versa for the other trials. In addition, to support learning and to allow solution quality anticipation (Czaczkes, Koch, et al. 2018), we also scented the paper overlays covering the runway leading to the feeder. Paper overlays were scented by storing them for at least one day in an airtight box containing a drop of essential oil on filter paper in a petridish. Finally, in addition to odours cueing sucrose molarity, visual cues were also provided. These consisted of printed and laminated pieces of paper (22 × 16.5 cm, figure 1B) displayed at the end of the runway, directly behind the sucrose droplet. Runways were scented on both outbound and inbound visits of the ant.

On the 9^th^ (test) visit, the odour and visual cue associated with either 1.5M or 0.1M were presented, while the sucrose solution provided was of intermediate (0.387M) quality, but also scented according to the runway scent. Runway scents in the test visit were varied systematically between ants, but each ant was confronted with only one of the two runway scents coupled with scented 0.387M sucrose.

Previous work has shown that *L. niger* foragers can form robust expectations of upcoming reward quality based on runway odour after 4 visits to each odour/quality combination (Czaczkes, Koch, et al. 2018; Wendt et al. 2019). Nonetheless, to ensure that learning had taken place, at the end of each training and test runs, we carried out a memory probe. The linear runway was replaced with a Y-maze (figure 1B), with two 10cm long arms and a 10cm long stem. The Y-maze stem was covered with an unscented paper overlay while one arm was covered with the 1.5M-associated odour overlay, and the other with the 0.1M-associated odour overlay. The matching visual cues were placed directly behind the relevant Y-maze arms. Trained ants were allowed to walk onto the Y-maze and their arm choice was noted. We used two decision lines to define arm choice – an initial decision line (fig. 1B, 2.5cm after the bifurcation) and a final decision line (7.5cm after the bifurcation). 91% of ants chose the side in the Y-maze which was covered in an odour previously associated to high molarity food and thus made a correct decision. Furthermore, on the test visit, ants deposited significantly more pheromone when presented with a high quality associated odour on the runway on their way to the food source compared to when the runway was impregnated with a low quality associated odour (online supplement figure S1B). Pheromone depositions towards the high quality odour increased with increasing experience with the food source during training, while they decreased for the low quality odour (online supplement figure S1A). This shows that they were able to associate a given odour to a food quality and could use this learned information on future foraging trips. After testing on the Y-maze, the ants were permanently removed from the colony.

## Statistical Analysis

Statistical analyses were carried out in R v. 3.5.0 (R Core Team 2016) using Generalized Linear Mixed Models (GLMMs) in the LME4 package (Bates et al. 2014) to analyse first interruption times, total drinking times and pheromone depositions data and Cumulative Link Mixed Models (CLMMs) in the ordinal package (Christensen 2015) to analyse food acceptance scores. CLMMs were used to analyse the acceptance data since we used an ordered factor with three levels (1 = full acceptance, 0.5 = partial acceptance, 0 = rejection).

As multiple ants were tested per colony, colony identity was added as a random effect to each model. GLMMs were tested for fit, dispersion and zero inflation using the DHARMa package (Hartig 2017). The model predictors and interactions were defined *a priori*, following Forstmeier and Schielzeth (2011). All p-values presented were corrected for multiple testing using the Benjamini–Hochberg method (Benjamini and Hochberg 1995). A total of 70 ants were tested – 34 with low quality associated cues and 36 with high quality associated cues.

### Food acceptance data

Model formula slightly differed depending on the experimental phase (training = visits 1 to 8, test = visit 9). Fixed factors used for statistical analysis of the training phase were “presented molarity” (1.5M or 0.1M) interacting with the “visit number” (1 to 8). Visit number was brought into the model as an interaction with presented molarity, because molarities were presented in an alternating order, always starting with low molarity on the first visit. Because individual ants were tested multiple times, we included AntID nested in colony as a random factor for statistical analyses of the training visits.

We used the following model formula for statistical analysis of the training visits:

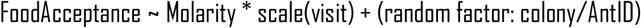

The fixed factor used for statistical analysis of the test visit was “high or low molarity associated odour cues” (odours were associated to 1.5M and 0.1M during the training phase).

This resulted in the following model formula for the test visit:

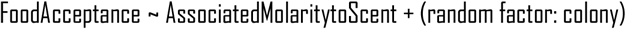

### First Interruption Times, Total Drinking Times & Pheromone Deposition Data

The total drinking times and pheromone deposition data were analysed using a GLMM with a Poisson distribution for: total drinking time during the test visit, and first interruption times and pheromone depositions for the training phase and the test visit. Total drinking times of the training phase were tested with a negative binomial distribution to receive a better model fit.

Model formula again slightly differed depending on the experimental phase (training = visits 1 to 8, test = visit 9). Fixed factors used for statistical analysis of the training phase were “presented molarity” (1.5M or 0.1M) interacting with the “visit number” (1 to 8). Visit number was brought into the model as an interaction with presented molarity, because molarities were presented in an alternating order, always starting with low molarity on the first visit. Because individual ants were tested multiple times, we included AntID nested in colony as a random factor for statistical analyses of the training visits.

We used the following model formulas for statistical analysis of the training visits:

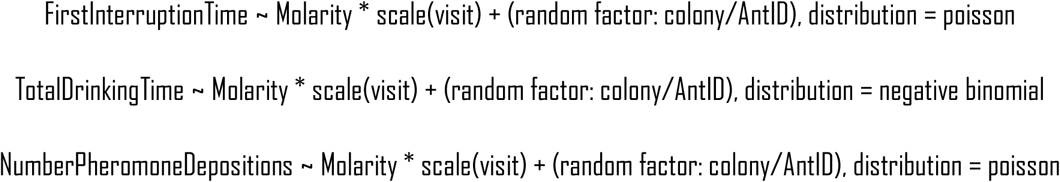

The fixed factors used for statistical analysis of the test visit were “high or low molarity associated odour cues” (odours were associated to 1.5M and 0.1M during the training phase) and the used odours (rosemary or lemon).

This resulted in the following model formulas for the test visit:

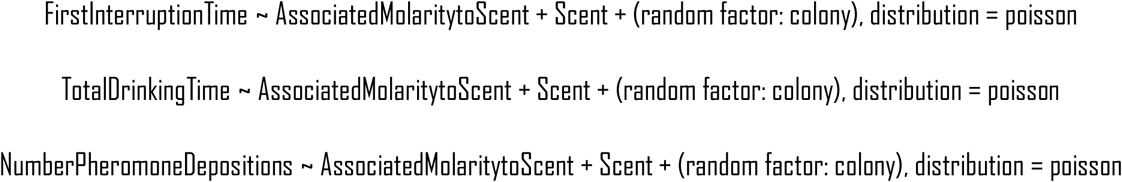

## Results

During training (visit 1 to 8), ants showed a higher total drinking time in seconds when confronted with the high molarity than when confronted with the low molarity (Estimate = -3.02, z = -41.81, p < 0.001, fig. 2A). In the test visit, the quality indicated on the runway and in the medium quality (0.387M) food strongly affected total drinking times. Drinking times were significantly higher when high-quality associated odours were present than when low-quality odours were present (median drinking time with high molarity cues: 44.63 seconds, low molarity cues 21.38 seconds, Estimate = -1.18, z = -4.47, p < 0.001, fig. 2B).

**Figure 2:**
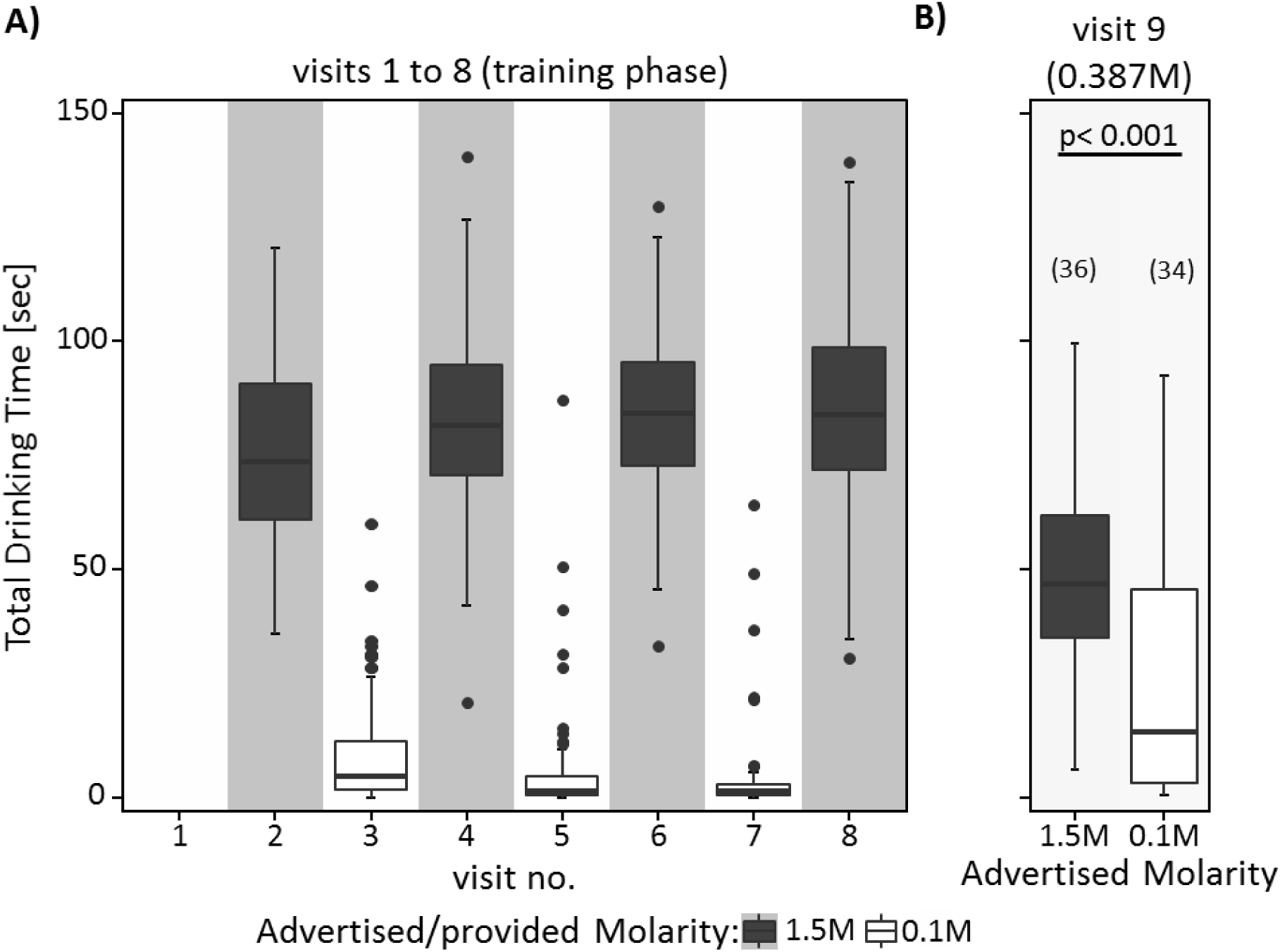
A) & B) Total Drinking Time in seconds for A) the eight training visits (visits 1-8) in which ants received 0.1M coupled with one scent and 1.5M coupled with another scent in an alternating order, always starting with 0.1M, B) the test visit (visit 9) in which ants always received 0.387M sucrose solution. Both the sucrose solution and the runway leading towards the food source were impregnated with one of the learned scents, triggering an expectation towards receiving either high or low molarities at the end of the runway. There is no data for total drinking time of the first visit displayed, because ants were sometimes disturbed when marking them, resulting in sometimes unclear feeding patterns. Shown are the median of total drinking time and the 75%/25% quantiles for each visit. Sample sizes for the 9^th^ visits of both experiments are displayed in parentheses of B).

Acceptance scores during training mirrored total drinking times, with ants showing a significantly higher food acceptance when confronted with the high molarity than when confronted with the low molarity (Estimate = -4.54, z = -4.97, p = <0.001, fig. 3A). However, food acceptance scores did not differ significantly in the test visit between the two advertised qualities (Estimate = -0.69, z = -1.51, p = 0.13, fig. 3B). However, there is a trend towards higher acceptance of the high-quality advertised food source. This is in contrast to the pattern found in Wendt et al. (2019), in which ants exposed to 1.5M-associated cues during the test visit showed significantly lower food acceptance towards the unscented 0.5M feeder than ants exposed to 0.25M-associated cues (CLMM: estimate= 1.07, z= 2.15, p= 0.03, figure 3C).

**Figure 3:**
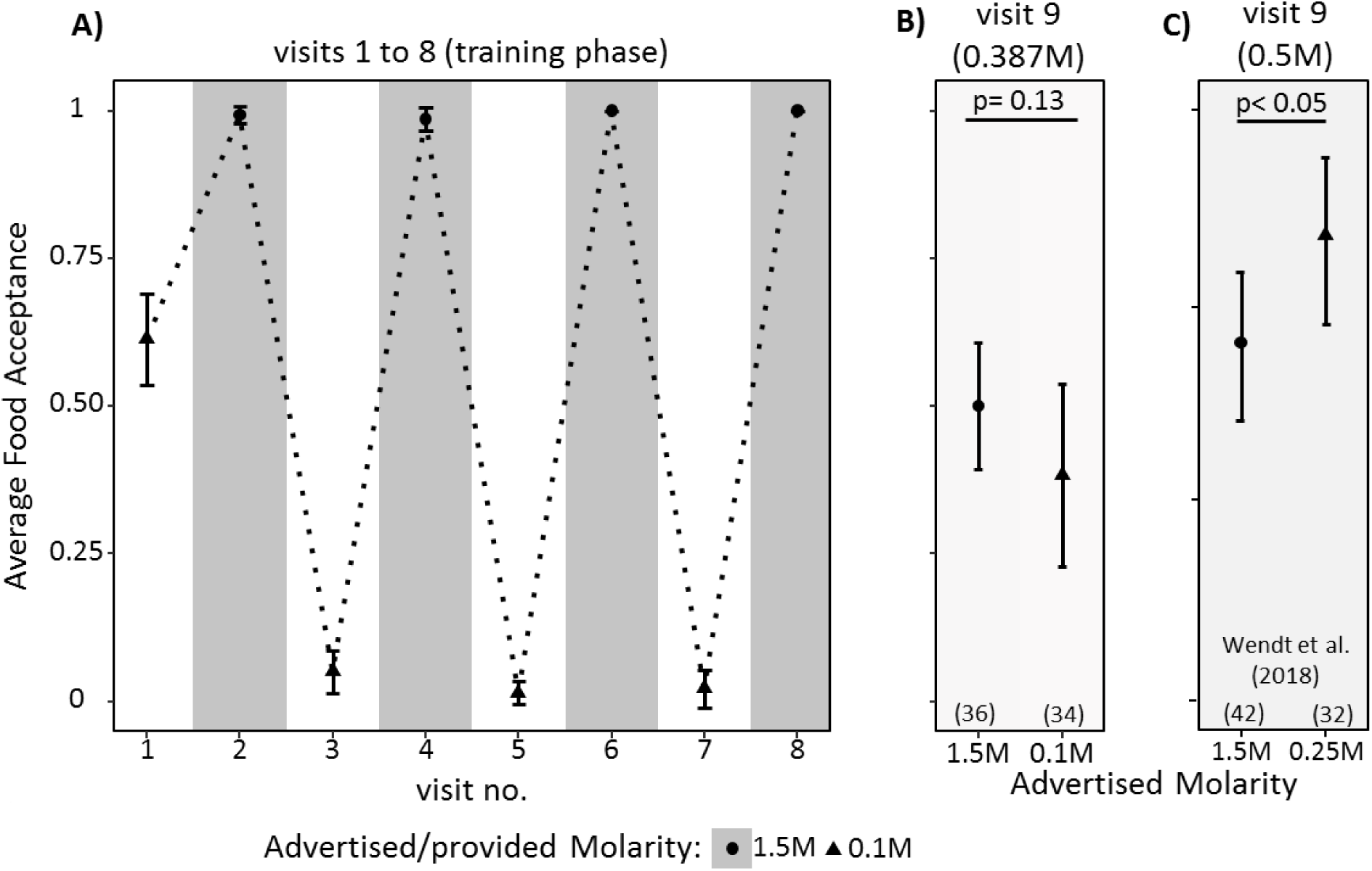
Average food acceptance for A) the eight training visits (visits 1-8) B) the test visit (visit 9) and C) the test visit (visit 9) of ants tested in Wendt et al. (2019) in which only the runway, but not the medium quality food (0.5M) was impregnated with learned odours. Shown are the mean food acceptance (points) and the 95% confidence intervals (error bars) for each visit. Sample sizes for the 9^th^ visits of both experiments are displayed in parentheses of B) and C).

First interruption times also mirrored acceptance scores and total drinking times during training, with higher first interruption times for the high quality food (Estimate = 0.53, z = 3.60, p < 0.001). In the test phase, there was a strong tendency towards ants showing lower first interruptions times for medium quality food advertised as high quality (Estimate = 0.71, z = 1.88, p = 0.06, figure 4B).

**Figure 4:**
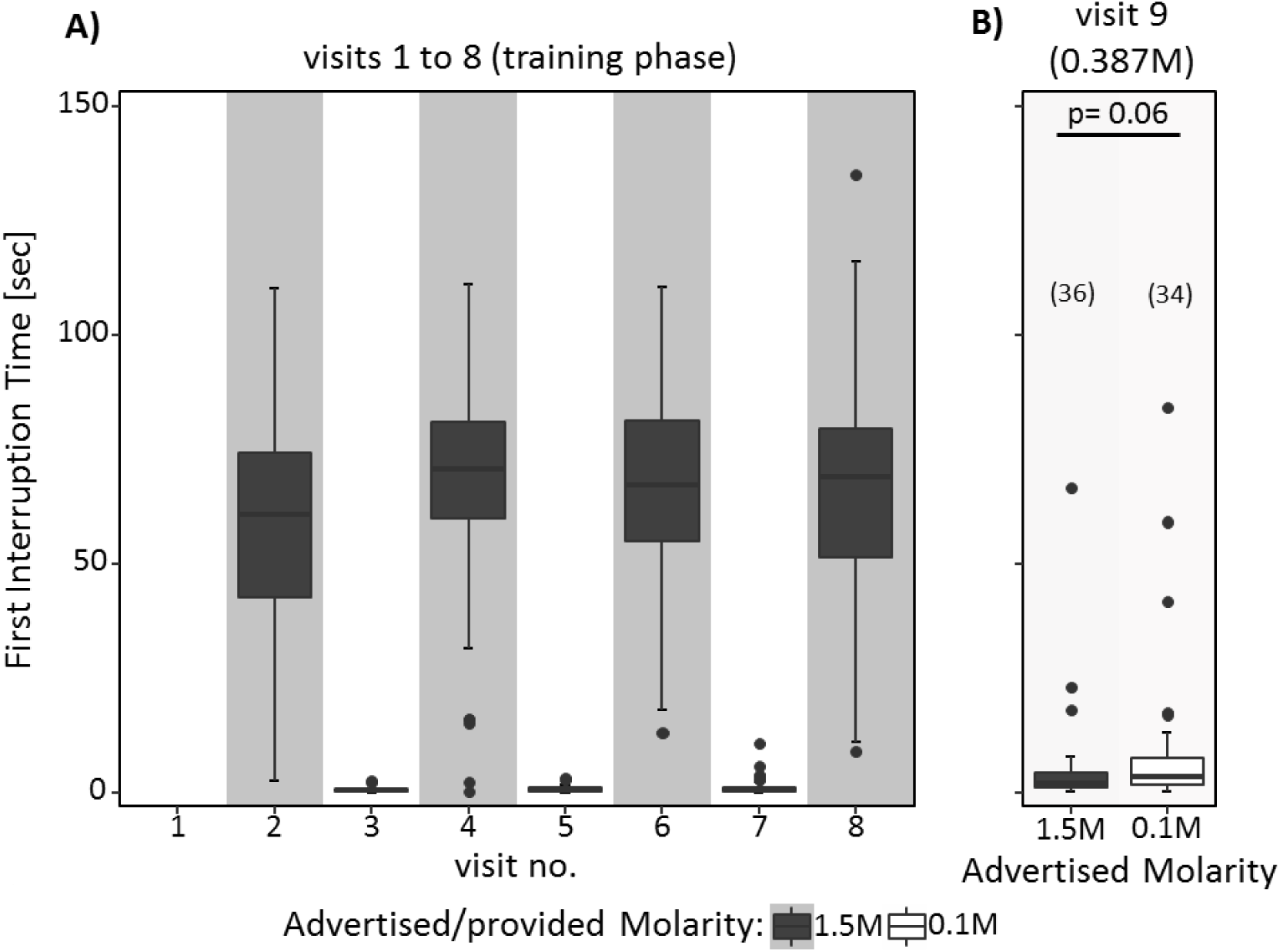
A) & B) First interruption time in seconds for A) the eight training visits and B) the test visit (visit 9). See figure legend 2 for details.

Finally, pheromone depositions when returning from the feeder to the nest also mirrored the other measured variables, with higher pheromone deposition for higher quality (Estimate = -1.58, z = -5.28, p <0.001, online supplement figure S2). On the test visit, advertised food quality affected pheromone deposition, with ants depositing more pheromone having consumed medium food advertised as high quality (Estimate = -3.19, z = -3.24, p < 0.01, fig. S2). However, note that ants deposited much less pheromone on the return from the test visit than on training visits (Estimate > 1.15, z > 9.37, P < 0.001). The data reported are similar to Wendt et al. (2019). Ants experiencing 1.5M-associated cues during the 9^th^ visit – provided only through runway scents towards the food, but not in the food – showed a significantly higher number of pheromone depositions on their return than ants exposed to 0.25M-associated cues (GLMM: estimate= -1.36, z= -5.50, p<0.001).

## Discussion

Ants spent more time feeding at medium quality (0.387M) sucrose solution when it was accompanied by a positively associated odour (previously associated to high molarity food) compared to a negatively associated odour (associated to low molarity food (figure 2B)). The number of pheromone depositions performed after feeding on medium quality food was also significantly higher when ants returned to the nest from 0.387M accompanied with high quality odour compared to low quality odour. Thus, ants reacted differently to food sources of identical sucrose solution, depending on the associative cue presented during consumption.

However, ants also showed some evidence of contrast effects in the first seconds after finding the food. The time until first feeding interruption at medium quality food was almost twice as long when ants expected low quality compared to expecting high quality food (p=0.06), suggesting that the medium quality food was perceived as better when ants expected to find poor food, and vice versa (fig. 4B). Finally, also as in Wendt et al. (2019), there was a significant difference in pheromone deposition depending on the ants’ expectations (online supplement figure S2). However, this experimental manipulation seems to interfere with pheromone laying, and the number of pheromone depositions is generally so low on the test visit that the difference does not seem biologically meaningful.

Our results should be considered in the context of our previous findings (Wendt et al. 2019). Previously, we performed a similar experiment, with a few modifications. Firstly, the food qualities used in the current experiment were different (0.1, 0.387, and 1.5M used here rather than 0.25, 0.5 and 1.5M) in order to balance the low and high quality food sources relative to medium quality. The generally lower food acceptance scores of the medium food in the current study were likely driven by this change. However, the critical difference is that in Wendt et al. (2019) the medium quality food was unscented, while in the current experiment it was scented. We believe it is this change which drives a reversed acceptance between the two experiments (compare figure 3B vs 3C). Wendt et al. (2019) showed clear contrast effects, where expectations caused an inversion in perception, so that high expectations caused an undervaluing of medium quality, and vice versa for low expectations. Here, with the minor addition of odour in the food, we reversed this pattern, resulting in an assimilation effect: if a label was present in the food indicating high quality during consumption, the perceived quality of the food increased. This assimilation effect can be very clearly seen in the total drinking time data as well (fig. 2B), and to a lesser extent in the pheromone deposition data. Hovland et al. (1957) argued that assimilation effects are likely to occur in humans when the expectation is not very different from the received option, whilst contrast effects are more likely to occur when the expectation is very different from the received option. Our results of this study together with those of Wendt et al. (2019) support this assumption in ants as well. The presence of an associated odour during consumption leads to a higher similarity between expectation and experience in the current experiment, in turn leading to assimilation rather than contrast effects which were shown in the previous study (Hovland, Sherif, and Harvey 1957).

We argue that this is directly analogous to the labelling effect described in humans. There, brand labels are based upon an accumulation of associative cues which have been linked to a label affecting perceived value (Levin, Schneider, and Gaeth 1998; Macklin 1996; Mao et al. 2013; van Osselaer and Janiszewski 2001). Just as humans prefer to purchase a brand with which they have previously had positive experiences over a novel brand (Russo, Medvec, and Meloy 1996), ants may also be affected by a familiar food label (associated odour) which previously offered positive (or negative) experiences, and may thus be more (or less) likely to “buy” a novel medium quality food source if it is presented with the familiar odour cue.

Our findings extend those of Oberhauser & Czaczkes (2018) who trained *Lasius niger* workers to a 1M food source presented along with either lemon or rosemary odour. After training, ants received a food source of identical quality, but presented with an unfamiliar odour. Ants showed significantly lower food acceptance towards the unfamiliar odour. There, as in this study, it is likely that the naturally value-neutral odour cue gained an associated value which affected value perception. Once the associated cue was missing, the reward lost part of its assigned value, leading to contrast effects, as also shown in Wendt et al. (2019). We propose that in the current experiment the odour on the runway and the taste of the food are playing different roles in the ant’s evaluation process: the odour is signaling what to expect, setting a reference point against which the measure of value obtained during feeding is contrasted. The taste is adding an associated value (positive or negative) during feeding, which is added to the objective sensory measure of food quality to form the complete measure of value obtained during feeding. This is then contrasted against the ant’s expectation.

Ants and bees can use odour information received inside the nest to find available food sources outside the nest (Grüter, Balbuena, and Farina 2008; Grüter and Farina 2009; Provecho and Josens 2009; Roces 1994; Saverschek and Roces 2011) and an unexpected odour can lead to lower acceptance of the same food source when it was previously presented with another odour (Lindauer 1949; Oberhauser and Czaczkes 2018). Associated odours thus strongly affect insect behavior and lead to different outcomes depending on where and when an odour is presented during a decision. Our results support the prediction that the presence of an associative cue during food consumption affects value perception, and that it can counteract expectations – even if the expectations and the associations are triggered by the same cue.

We demonstrated an assimilation effect driven by labels in ants, mirroring effects found in humans. This adds to prior studies showing parallel cognitive preferences between humans and insects, including decoy effects (Sasaki and Pratt 2011; Tan et al. 2014, 2015), risk aversion (De Agrò, Grimwade, and Czaczkes 2019; Shafir et al. 1999; Shapiro 2000; Waddington, Allen, and Heinrich 1981), discounting (Cheng et al. 2002; Wendt and Czaczkes 2017) and expectation-driven valuation (Bitterman 1976; Couvillon and Bitterman 1984; Wendt et al. 2019), suggesting that insights into human behaviour can, in part, be transferred to insects. Insect-based comparative psychology studies allow much tighter control over experimental subjects and conditions, offering stringent tests of basic insights from human psychology, and the experimental flexibility to test hypotheses untestable on human subjects. We hope that our work inspires consumer psychologists and behavioural economists to consider insects as a viable model system in which to test their underlying assumptions and thinking, in order to gain deeper insights in both human and animal behaviour.

## Acknowledgements

We thank Kim S. Strunk for helpful comments on an earlier version of this manuscript. We also thank the DFG (Deutsche Forschungsgemeinschaft) which funded SW and TJC with an Emmy Noether grant to TJC, grant number CZ 237/1-1.

## Author contributions

SW performed the experiments and analysed the data. SW and TJC designed the study, interpreted the data and wrote the manuscript. All authors gave final approval for publication and agree to be held accountable for the content therein.

## Ethical approval

All animal treatment guidelines applicable to ants under German law have been followed.

