## Supplementary material for "Labelling effect in insects: cue associations influence perceived food value in ants": online supplement

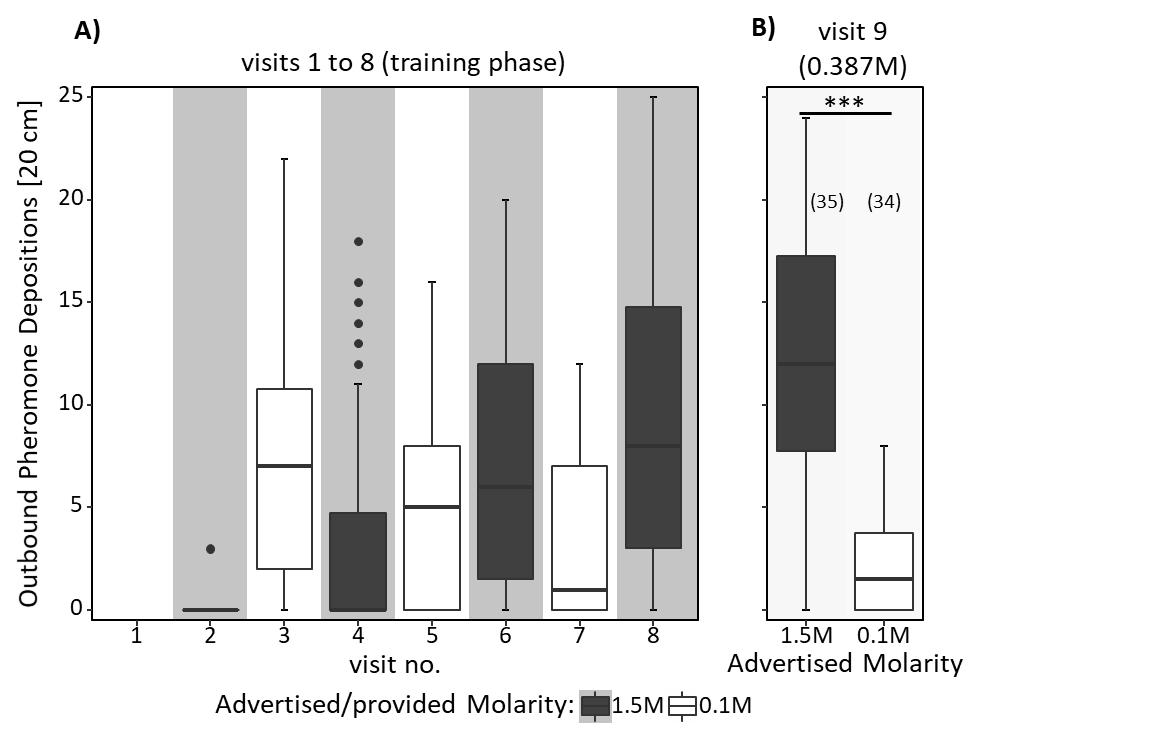

Figure S1: A) & B) Outbound Pheromone Depositions [20 cm] (to the Food Source) for A) the eight training visits (visits 1-8) in which ants received 0.1M coupled with one scent and 1.5M coupled with another scent in an alternating order, always starting with 0.1M, B) the test visit (visit 9) in which ants always received 0.387M sucrose solution. Both the sucrose solution and the runway leading towards the food source were impregnated with one of the learned scents, triggering an expectation towards receiving either high or low molarities at the end of the runway. Shown are the median number of pheromone depositions on the measured 20cm track on the way to the food source and the 75%/25% quantiles for each visit.

During training (visit 1 to 8), ants deposited significantly more pheromone on the measured 20 cm track on the way to the food source when confronted with the high molarity than when confronted with the low molarity (Estimate = 0.52, z = 4.65, p < 0.001). The number of visit also had a significant effect on the number of deposited pheromone with pheromone depositions towards high molarity scent generally increasing and pheromone depositions towards low molarity scents generally decreasing over time (Estimate = 1.14, z = 13.64, p < 0.001). Ants which expected high molarity on the 9^th^ visit (test visit) deposited significantly more pheromone than ants which expected to find low molarity food at the end of the runway (Estimate = -1.80, z = -8.85, p < 0.001). Number of outbound pheromone depositions were also significantly higher for ants confronted with the high molarity scent on the ninth visit compared to the training phase (visit 2 vs 9: Estimate = 0.46, z = 11.14, p < 0.001, visit 4 vs 9: Estimate = 0.57, z = 13.96, p < 0.001, visit 6 vs 9: Estimate = 0.59, z = 14.5, p < 0.001, visit 8 vs 9: Estimate = 0.58, z = 14.36, p < 0.001).

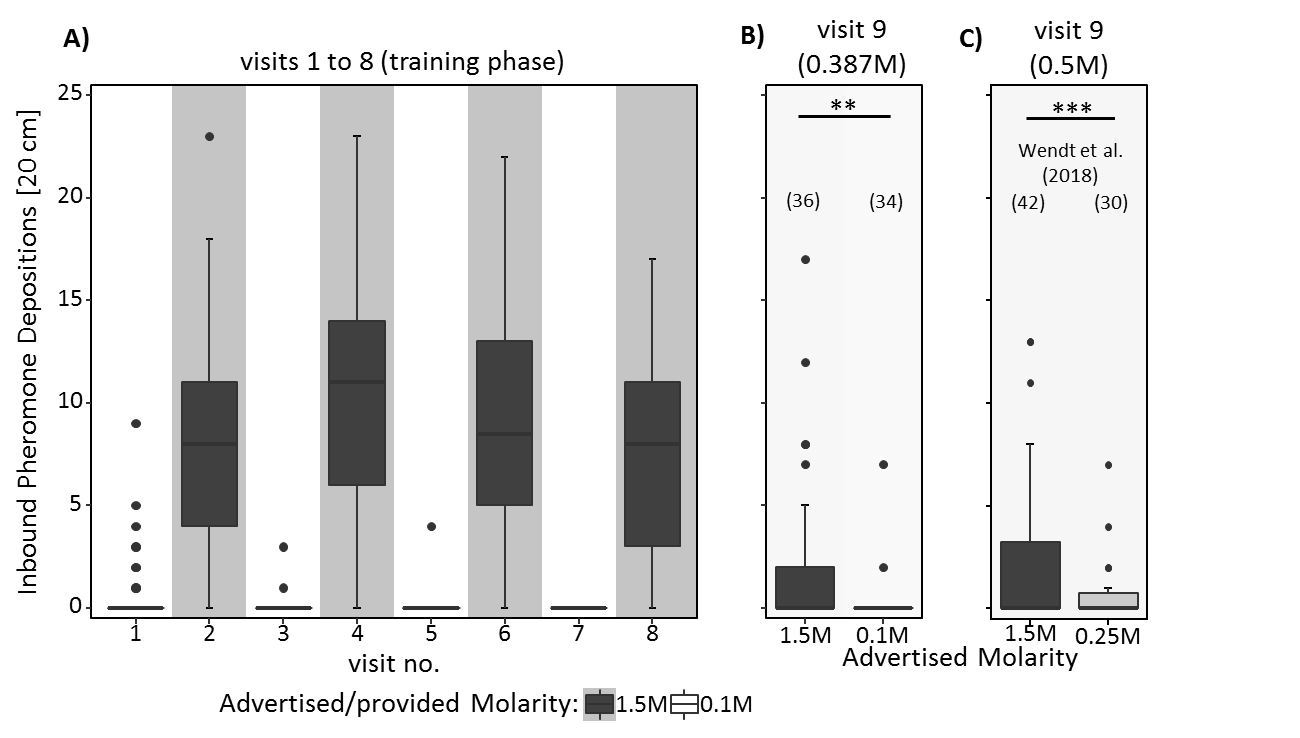

Fig. S2: Pheromone depositions towards the nest [20 cm] (inbound pheromone depositions) of the data presented in (Wendt et al. 2018) for A) the eight training visits (visits 1-8) in which ants received 0.25M coupled with one scent and 1.5M coupled with another scent in an alternating order, always starting with 0.25M, and B) the test visit (visit 9) of ants tested in Wendt et al. (2018) in which only the runway, but not the medium quality food (0.5M) was impregnated with learned odours. Shown are the median number of pheromone depositions on the measured 20cm track on the way back to the nest and the 75%/25% quantiles for each visit.
